# Accurate fetal variant calling in the presence of maternal cell contamination

**DOI:** 10.1101/552414

**Authors:** Elena Nabieva, Satyarth Mishra Sharma, Yermek Kapushev, Sofya K. Garushyants, Anna V. Fedotova, Viktoria N. Moskalenko, Tatyana Serebrenikova, Eugene Glazyrina, Ilya V. Kanivets, Denis V. Pyankov, Tatyana V. Neretina, Maria D. Logacheva, Georgii A. Bazykin, Dmitry Yarotsky

**Affiliations:** Skolkovo Institute of Science and Technology, Skolkovo, Russia; Institute for Information Transmission Problems (Kharkevich Institute), Russian Academy of Sciences, Moscow, Russia; Lomonosov Moscow State University, Moscow, Russia; Progen LLC, Moscow, Russia; Genomed LLC, Moscow, Russia

**Keywords:** maternal cell contamination, prenatal sequencing, variant calling, fetal exome sequencing

## Abstract

High-throughput sequencing of fetal DNA is a promising and increasingly common method for the discovery of all (or all coding) genetic variants in the fetus, either as part of prenatal screening or diagnosis, or for genetic diagnosis of spontaneous abortions. In many cases, the fetal DNA (from chorionic villi, amniotic fluid, or abortive tissue) can be contaminated with maternal cells, resulting in the mixture of fetal and maternal DNA. This maternal cell contamination (MCC) undermines the assumption, made by traditional variant callers, that each allele in a heterozygous site is covered, on average, by 50% of the reads, and therefore can lead to erroneous genotype calls. We present a panel of methods for reducing the genotyping error in the presence of MCC. All methods start with the output of GATK HaplotypeCaller on the sequencing data for the (contaminated) fetal sample and both of its parents, and additionally rely on information about the MCC fraction (which itself is readily estimated from the high-throughput sequencing data). The first of these methods uses a Bayesian probabilistic model to correct the fetal genotype calls produced by MCC-unaware HaplotypeCaller. The other two methods “learn” the genotype-correction model from examples. We use simulated contaminated fetal data to train and test the models. Using the test sets, we show that all three methods lead to substantially improved accuracy when compared with the original MCC-unaware HaplotypeCaller calls. We then apply the best-performing method to three chorionic villus samples from spontaneously terminated pregnancies.

**Code and training data availability:** https://github.com/bazykinlab/ML-maternal-cell-contamination

## Introduction

High-throughput sequencing of fetal DNA is increasingly being used in academic and clinical settings. It is a powerful tool with the potential for use in prenatal diagnosis based on chorionic villus or amniotic fluid sampling [1], or in the analysis of chorionic villi or products of conception for genetic diagnosis of an unsuccessful pregnancy. In prenatal diagnosis, whole-genome or whole-exome sequencing can discover novel clinically significant variants that are not present in SNP arrays or gene panels, resulting in higher diagnostic yield [2]. Prenatal sequencing can inform prenatal and postnatal care and counseling, and may lead to prenatal therapeutic interventions [2]. In standard practice, the DNA of both parents is sequenced together with fetal DNA (“trio sequencing”) in order to establish patterns of inheritance and inform variant prioritization and interpretation. A technical difficulty that may arise in the analysis of fetal DNA is the contamination of the fetal sample with maternal cells. The prevalence of such *maternal cell contamination* (MCC) can be significant, depending on the experimental technique and quality of the sample; for example, one study reported 9.1% of amniotic fluid samples as having detectable MCC [3], while another found MCC fraction *>* 5% in as many as 26% of amniotic fluid samples under some practices [4]. High-level contamination (over 20%) was detected by one study [5] in a small, but non-zero number of samples (0.3% of cultured amniotic fluid samples and 1.3% of cultured chorionic villi samples; it must be noted that cultured amniotic fluid samples generally have less MCC than direct samples). In traditional prenatal analysis, such as that aimed at detecting chromosomal aberrations, maternal cell contamination is assayed by special tests, such as the Short Tandem Repeat analysis, and, if detected at a sufficient level, may nullify the analysis [6].

Meanwhile, standard variant calling software that is used to analyze next-generation sequencing data relies on the expectation that each allele is represented by half of the reads. MCC disrupts this assumption, leading to errors in variant calling. In this work, we propose and evaluate computational methods for reducing the MCC-caused error. All of these methods begin with variants called in the fetal specimen, the mother, and the father by a standard variant-calling pipeline and then “correct” the results from the fetal specimen. The first method uses Bayesian estimator to decide on the “true” fetal genotype based on the called genotypes of the trio. The other methods eschew making assumptions about the best way to uncover the “true” fetal genotype from the maternally-contaminated observed specimen data and instead solve this problem using machine learning. We train these methods on synthetic “mother-father-fetus” trios generated from real family trios by adding specified numbers of maternal reads to the child sample. We use these synthetic trios, where the child’s genotype is known, to demonstrate that MCC correction significantly improves the accuracy of variant calling compared to contamination-naive calling, especially for higher fractions of MCC. As an intermediate technical step, we present a simple heuristic algorithm for estimating maternal cell contamination fraction in the fetal sample. We then apply the trained model to real sequencing data from a miscarried fetus with 40% estimated MCC, changing the fetal calls for a substantial number of SNPs.

## Materials and methods

In the rest of the paper, we use the term *specimen* to denote the obtained fetal sample which may be contaminated with maternal cells. In practice, it can be a chorionic villus or amniotic fluid sample or an abortus sample.

We assume that DNA samples are available for all three members of the trio. We expect the trio to be sequenced (we gear our work towards and test it on Illumina whole-exome sequencing, currently the most widely used protocol in biomedical practice) and the reads mapped and variant-called according to a standard bioinformatics pipeline, e.g., one involving the Genome Analysis Toolkit (GATK, [7]), although our approach can work with other calling pipelines too, as we show in the section *Strelka2 genotype correction*. We expect that the called genotypes are available along with accompanying quality information, such as allelic depths, genotype likelihoods, and so on. The VCF file produced by the pipeline is the starting point for all our analyses (see Fig. 1).

**Fig 1.**
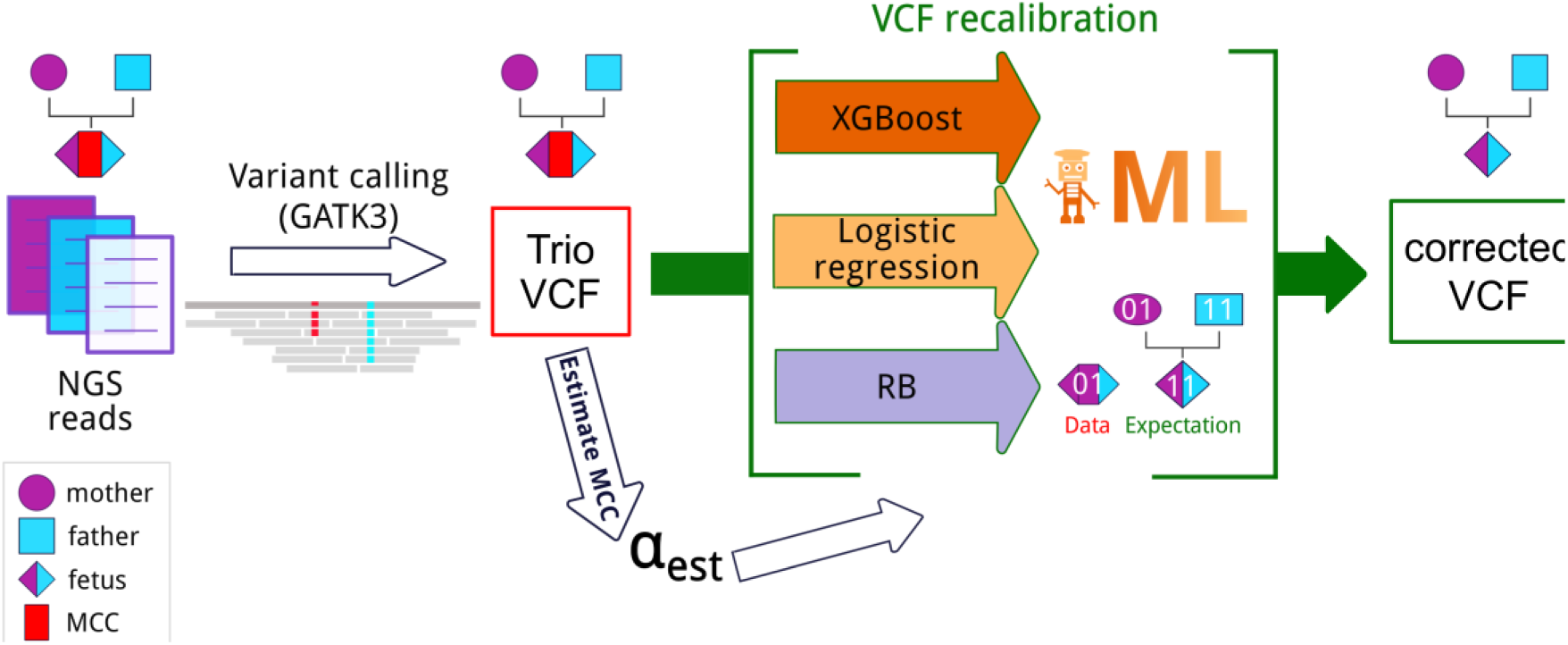
Pipeline for accurate fetal variant calling in the presence of maternal cell contamination (MCC). NGS reads for each sample in the trio are mapped to the reference genome, and then variants are called with GATK v3. In the presence of MCC, the resulting VCF file contains incorrect calls for the fetus. To overcome this, we estimate MCC from this VCF and utilize the estimated value to correct the genotype calls for the fetus. Three different approaches for genotype correction were utilized (see Materials and Methods section): a restricted Bayesian method (RB) and two machine learning-based approaches, namely, logistic regression and Gradient Boosted Decision Trees (XGBoost). ML - machine learning.

### Estimating the contamination fraction

We define the maternal cell contamination (MCC) fraction, which we denote by *α* ∈ [0,1], to be the share of maternal DNA in the DNA of the fetal specimen. MCC accounts for the discrepancy between the results of variant calling on the contaminated specimen sample and the true fetal genotype. The MCC-aware genotype correction methods presented below all rely on knowing the value of *α*, and it is vital to be able to estimate it accurately.

To estimate *α* in a fetal sample, we look at the results of variant calling by GATK HaplotypeCaller [7] and consider positions in the genome where the mother and the father are homozygous for different alleles (i.e., one of the parents is homozygous for the reference allele, and the other for the alternative allele); we only consider biallelic sites. The fetus then should be heterozygous at these sites, with equal read coverage for the two parental alleles.

In the presence of MCC, however, the mother’s allele coverage fraction *m* should be higher, namely 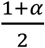. Then, we can use the value of *m* at the position obtained from the VCF file to compute *α* as 2*m* − 1. Since the actual coverage fluctuates, we average this ratio over all relevant sites to get the MCC fraction estimate. Let *m*_*i*_ be the fraction of maternal reads at site *i*. Then we compute 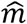 to be the average of *m*_*i*_ over all sites *i* where the mother and the father are homozygous for different alleles, and estimate 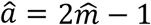. Only calls that pass basic filtering criteria (GQ *>* 30 in both parents, depth at site *>* 10 in all three samples) are considered in the calculation.

We observe that estimating *α* when the mother is homozygous for the reference allele (“mo00”) systematically gives larger values than when the mother is homozygous for the alternative allele (“mo11”). We attribute this phenomenon to reference bias [8]. We, therefore, estimate *α* separately over the two alternatives (“fa00_mo11” and “fa11_mo00”) and compute the average of the two estimates as the final result. We found this aggregated estimate to be more accurate than either alternative. Using simulated data (see section Contaminated trio generation), we found that the MCC estimation error made by this method does not exceed 2% and in the vast majority of the cases is below 1% (Fig. S6).

While there are methods for estimating sample contamination that are geared towards detecting contamination during sequencing with unrelated samples and make use of population allele frequencies (VerifyBamID [9] and the CalculateContamination module of the cancer-sequencing workflow in Genome Analysis Toolkit v.4 (https://software.broadinstitute.org/gatk/), they are not well suited for the particular case of maternal contamination; in fact, the authors of VerifyBamID predicted that their method would underestimate contamination with maternal sample by half [9]. Our evaluation of these methods on simulated contaminated data generally agrees with this prediction, although we find it to be something of an underestimate itself, especially for large MCC fractions (Fig. S6).

### Restricted Bayesian genotype correction

Our first method uses an explicit probabilistic model and the Bayesian approach. Let AD0, AD1 denote the allelic depths of the reference and alternative variants in the mother-fetus mixture, and cGT,mGT,fGT denote the genotypes of child, mother and father, respectively. The joint probability of these variables is factorized as

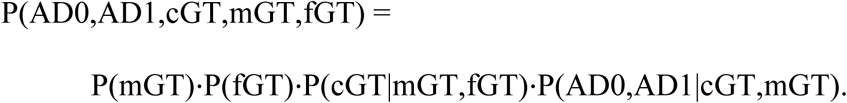

Here, the P(mGT) and P(fGT) are prior probabilities of mother’s and father’s genotypes, P(cGT | mGT,fGT) is the conditional probability describing Mendelian inheritance, and P(AD0, AD1|cGT,mGT) is the conditional probability describing observations of reads in the mixture for given genotypes of mother and child. The prior probabilities P(mGT) and P(fGT) are available from mother’s and father’s VCF files, while the conditional probabilities are given by explicit formulas. From this representation, we can find the conditional probabilities P(cGT|AD0,AD1) and hence determine the most likely genotype of the child given the allelic depths; see Section S3 for details.

Unlike a full-scale variant caller that has access to individual read data, we model all the read and alignment errors with a uniform error rate ε (see Section S3). We observe that setting ε to a small but non-zero value does lead to an increase in accuracy (Fig. S2).

If contamination α = 0, the predictions of the child’s genotype by the above Bayesian model should agree with the prediction of the standard pipeline applied to the mixture, but in reality we observe some discrepancy between the two (Fig. S1). This discrepancy is due in part to the Bayesian method’s lack of access to the individual read data, but also to differences between the Bayesian model and the variant-calling algorithm. For example, the Bayesian model takes into account Mendelian inheritance, while the GATK pipeline leaves it until subsequent Genotype Refinement steps [10],[11].

As the purpose of this work is not to replace existing variant callers, but to correct the errors they make due to maternal cell contamination, we focus on the scenarios where the standard variant calling pipeline likely makes a mistake, and replace the original call by the Bayesian prediction in these cases only. Specifically, we observe that the disagreement between the child and fetal specimen variants is in the vast majority of cases due to the mislabeling of homozygous positions in the fetus as heterozygous in the contaminated fetal specimen (Fig. S2). This happens when the maternal genotype, and, accordingly, the mixture of fetal and maternal reads, is heterozygous. Following this observation, we define the *Restricted Bayesian* model as follows: if both maternal and contaminated specimen’s genotypes (as found by the standard variant caller) are heterozygous, then the child’s genotype is recomputed using the above Bayesian model, otherwise it is left intact. This restricted Bayesian model performs well compared both to the unrestricted Bayesian model and to using the uncorrected predictions (Fig. S3)

### Machine learning-based genotype correction

In this approach, the predictive model for genotype readjustment is trained using a machine learning algorithm. The input to the model consists of two components: the estimated fraction of maternal DNA in the fetal specimen *α*_*glob*_, and the vector of features characterizing the variants called in the fetal specimen and the parents at a particular position in the genome (practically, this is the line in the VCF file for that position; it includes the genotype likelihoods, genotype qualities, allele-wise and total read depths, and the called genotype for all three samples). The output of the model is the corrected fetal genotype at this position in the genome. The fields describing the variant (genotype likelihood, read depth, etc.) need to be the same in the training VCFs and the VCF to be genotype-corrected; therefore, the same or closely related variant callers should be used to produce all VCFs. To train and test the model, we simulate “virtual specimens” from a number of publicly available father-mother-child trios by randomly mixing mother and child reads at various MCC fractions, as described in *Contaminated trio generation*, and then use the genotype calls for the pure fetal sample as the “ground truth” in training. We considered two machine learning algorithms: logistic regression (as implemented in scikit-learn (http://scikit-learn.org/) and Gradient Boosted Decision Trees (XGBoost implementation [12]). The inputs to the models are read from a VCF file and standardized, with categorical features being one-hot encoded (split into binary columns for each unique value the feature can take). L2 regularization is applied to both algorithms, along with a minimum loss reduction threshold for decision tree partitions and early stopping based on performance on a validation set used to prevent XGBoost from overfitting. We found both models to be robust to choice of hyperparameters, and while these can be further tuned for a particular dataset, the performance gains are marginal and do not warrant the loss in generalizability it incurs.

### Contaminated trio generation

We obtained testing and training datasets by using publicly available exome reads from father-mother-child trios (“real-world trios”) to produce “virtual specimen trios” with predetermined maternal contamination fraction *α*. Namely, we mixed randomly selected reads from the “real-world child” and the “real-world mother” in such a proportion that the fraction of maternal reads would be *α*, and thus obtained “virtual specimen” reads. We used the remaining reads from the “real-world mother” to create the “virtual specimen mother,” while the “virtual specimen father’s” reads were identical to the original “real-world father’s” reads (we split the mother’s reads into non-overlapping “specimen contamination” and “mother” subsets to avoid using the same information twice). For each “virtual specimen” trio, we also generated a corresponding “pure child” trio, which was identical to the “virtual specimen” trio except that no mother’s reads were added to the child’s reads. We kept the mother’s reads and the child’s reads identical both in the “real world” trio and the “virtual specimen trio” to keep the amount of information available to the caller in both settings identical and the comparison as fair as possible.

We performed this procedure for four “real-world” trios with publicly available exome data: the Ashkenazim trio HG002_NA24385 from the Genome in the Bottle project [13] (“AJT”), the YRI_NA19240 trio from the 1000 Genomes project [14] (“YRI”), the CHD trio [15] (“CHD”), and the Corpas family daughter trio [16] (“Corpas”). Since the child in the “real-world” trios was either a living individual (AJT, YRI, Corpas) or stillborn (CHD), there was no maternal cell contamination in the “real-world” reads. If the data were available only as a bam or cram file, we first obtained raw reads using the bam2FastQ program of the bamUtil package [17]. The CHD trio had a large number of read duplicates, which distorted the contamination fraction, so we deduplicated the aligned reads using the biobambam package (https://www.sanger.ac.uk/science/tools/biobambam), and used the reads from the deduplicated file to generate the “contaminated” trio.

For each “real-world trio”, we repeated this procedure with α = 0.01, 0.03, 0.05, 0.1, 0.15, 0.2, 0.25, 0.3, 0.35, 0.4, 0.45, 0.5. We aligned the “virtual specimen” and the “pure child” trio reads to the GRCh38DH reference genome following the 1000 Genomes pipeline [18]. Variants were called using Genome Analysis Toolkit 3.8 HaplotypeCaller with the option -dontUseSoftClippedBases and restricting the sequence considered to Gencode v.24 protein-coding exons.

Table 1 shows for each trio, the number of called variants and the coverage. Fig. S1 shows the distribution of virtual specimen genotypes and its variation with MCC.

**Table 1.**
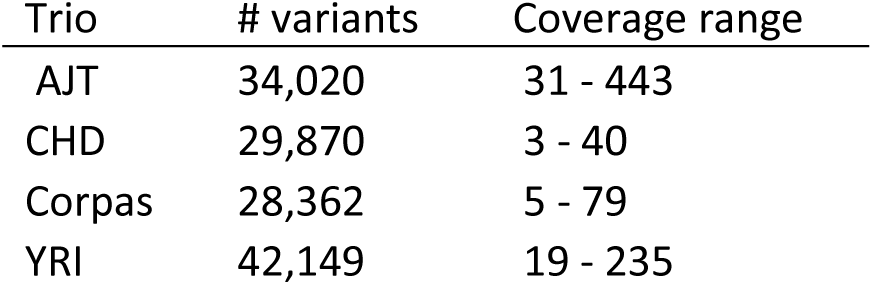
The number of variants and range of coverage for each synthetic trio. The number of variants is given for MCC fraction 0.01, and is reduced by up to 5% as MCC is increased to 0.5. The edges of the coverage range denote the fifth and ninety fifth percentile values for the virtual specimen variant read depth in the 0.01 MCC trio.

| Trio | # variants | Coverage range |
| --- | --- | --- |
| AJT | 34,020 | 31 - 443 |
| CHD | 29,870 | 3 - 40 |
| Corpas | 28,362 | 5 - 79 |
| YRI | 42,149 | 19 - 235 |

### Miscarriage samples

We analyzed DNA from spontaneously miscarried euploid abortuses. Internal Review Board approval was obtained from the Ethics Committee of the Institute of Information Transmission Problems (document 11616-2116/726, 27/11/2017) and the Institutional Review Board of the Skolkovo Institute of Science and Technology (20/06/2019). Families who suffered a miscarriage and requested chromosomal microarray analysis (CMA) of the embryo were offered the choice to participate additionally in the whole-exome sequencing study. Informed consent was obtained from the parents. Only cases without CMA-detected copy-number anomalies were considered for the WES study. Chorionic villi were used for the abortus DNA sample, while blood was drawn from both parents for sequencing. Parenthood and the presence of fetal DNA in the chorionic villus sample were verified using short tandem repeat (STR) analysis with the COrDIS Plus system (GORDIZ, Moscow, Russian Federation). Libraries were prepared using the TruSeq DNA Library Prep for Enrichment kit (Illumina, San Diego, CA, USA). Exome capture was performed using the xGen Exome Research Panel v.1.0 (Integrated DNA Technologies, Coralville, IA, USA). Sequencing was performed on the HiSeq4000 instrument (Illumina, San Diego, CA, USA) in paired-end mode. The reads were aligned to the hg19 reference genome. Calling was restricted to the Gencode v.2.7 protein-coding exons with 50-nt flanks. Otherwise, the mapping and calling procedure was the same as for the “virtual specimen” trios (see *Contaminated trio generation*).

## Results

We compared the genotypes corrected by each of the MCC-aware methods on the “virtual specimen trios” to the genotypes produced by the GATK HaplotypeCaller on the corresponding “pure child” trios, the latter serving as the ground truth. As a baseline, we also included the comparison to initial genotypes called by the GATK HaplotypeCaller on the “virtual specimen” trios (“no genotype correction”). To control for possible artifacts that may result from experimental differences in the original sequencing of the real-world trios, including differences in coverage, we present the results separately for the “virtual specimen” trios generated from each original family. The machine learning-based models were trained with a “leave-one-out” strategy, where every trio excluding the test trio was used for training. There are some input features that are strongly correlated, for example the genotypes and their associated likelihood scores. While this may present a challenge for the linear logistic regression model, gradient boosted decision trees are robust to multicollinearity by design.

Fig. 2 summarizes the results, with the accuracy achieved by each method plotted against the contamination fraction. All three individual methods fare well, reducing the number of miscalled positions by 40-80% over the baseline “no genotype correction” approach. Both machine learning approaches generally outperform the restricted Bayesian method and overcome some of its problems. XGBoost is the best performer. Restricting analysis just to indels similarly shows improvement in accuracy from genotype correction, even when training on non-indel variants (Fig. S7).

**Fig 2.**
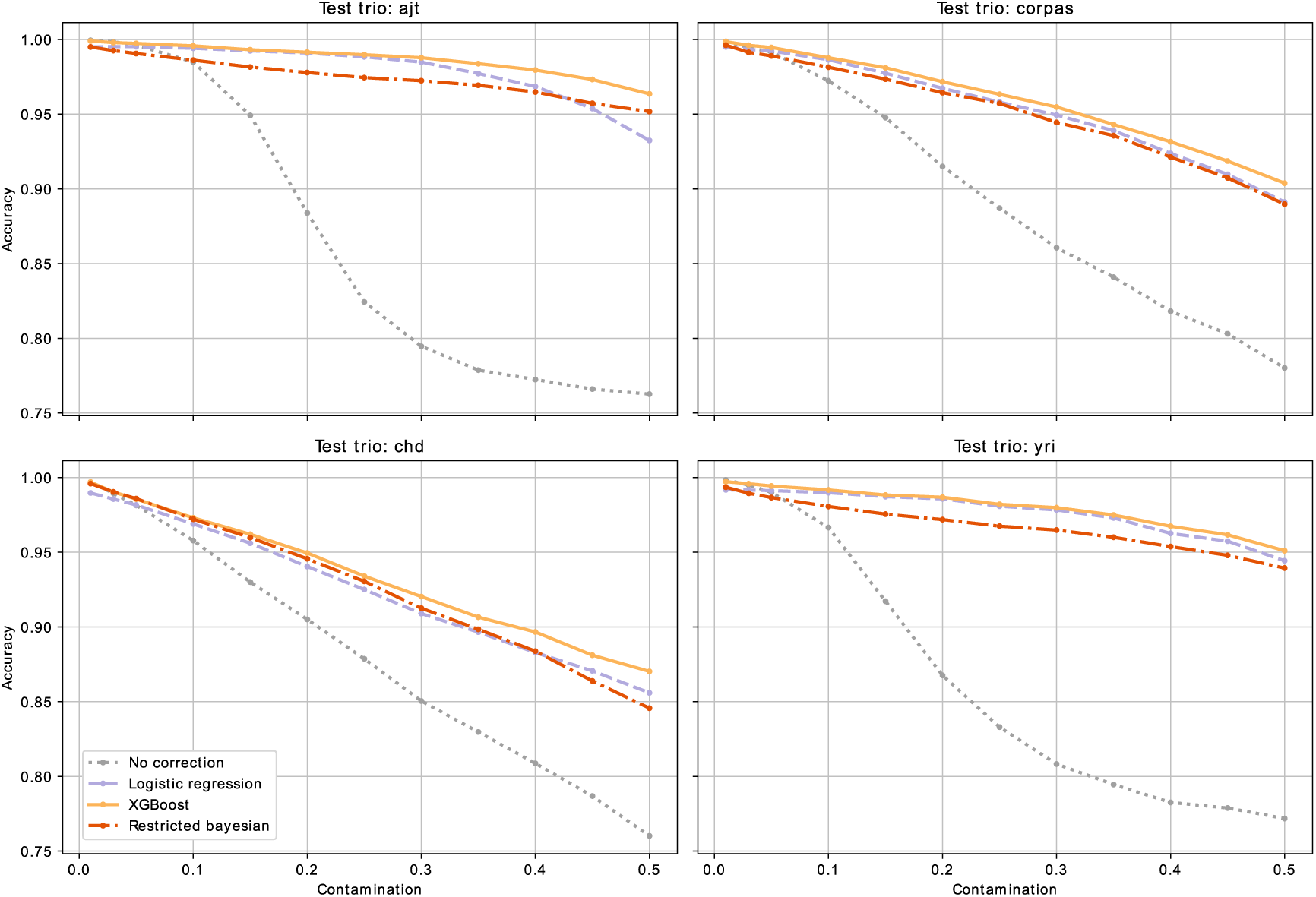
Accuracy of the genotype correction methods at various MCC fractions. Machine learning methods were trained with a “leave one out” strategy. Each curve consists of twelve data points corresponding to accuracy of the method applied to “contaminated specimens” at various MCCs (the contamination values used are as given in section Contaminated trio generation). The accuracy is calculated on the intersection of the calls made on the virtual fetal specimen and the calls made on the real-world child.

Fig. S5 visualizes the accuracy of XGBoost as compared to no correction on a per-genotype basis. Note that in our two lowest-read-coverage trios, it seems that correcting the genotype with XGBoost results in a lower accuracy for heterozygous variants: the accuracy on uncorrected heterozygous variants is higher at the cost of mislabelling a significant number of homozygous variants as heterozygous.

We also compared the running times of the methods for both training and testing (Table 2). In terms of speed, the restricted Bayesian method, which requires no training, is the clear winner. The machine learning methods take on the order of minutes to train. Once the models are trained, the times to genotype-correct a specimen are comparable.

**Table 2.**
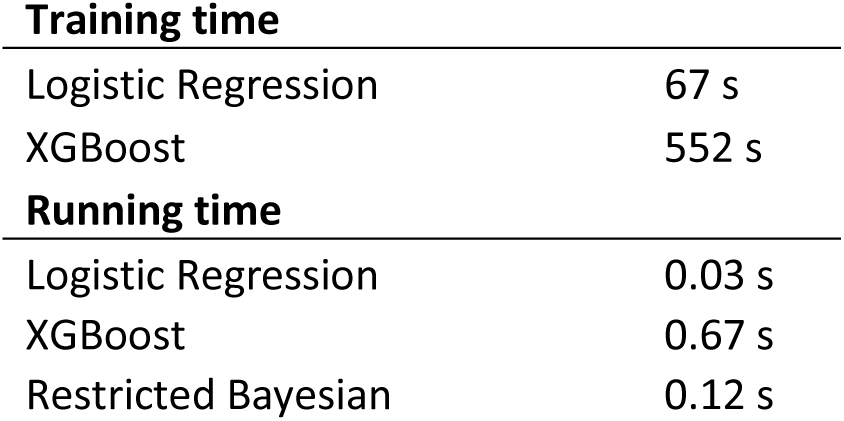
Training and running times of our methods. Training includes training a model on the entire synthetic dataset. Running involves genotype-correcting a single VCF file (96,000 variants) with a pre-trained model. The benchmarks were run on an Intel i5-7200U CPU @ 2.5 GHz.

### Factors affecting the quality of machine learning-based genotype correction

As noted earlier, the training dataset needs to be sufficiently rich for machine learning to succeed at genotype correction. To demonstrate this, the two machine learning algorithms were trained and tested on different pairs of families (a ‘one versus one’ approach), at all contamination fractions. The results are summarized in Fig S3. While the methods almost always lead to an improvement in accuracy, the gains are much more modest than when training with a ‘leave-one-out’ approach (Fig 2), due to overfitting to the features of a particular trio. This is particularly noticeable when correcting high-coverage trios with a model trained on a low-coverage trio. To combat overfitting, we ultimately adopted the approach where we train on all available trios except for the test one and added sufficient regularization to our models.

The trained classifiers can tell us about the discriminative power of our input features. Table 3 shows the top 10 features ordered by discriminative power for both classifiers. It is interesting to note that the degree of contamination is not found among the top features for logistic regression. Additionally, the father’s features are prominently towards the bottom of the table for logistic regression, yet are included in the top 10 for XGBoost, suggesting some complex relationship that the linear model does not capture. XGBoost likely considers the heterozygosity of the specimen as a starting point while constructing decision trees (which is consistent with the nature of genotyping errors, see Fig. S2), but then prefers a combination of genotypic likelihood scores and allelic depths while proceeding.

**Table 3.**
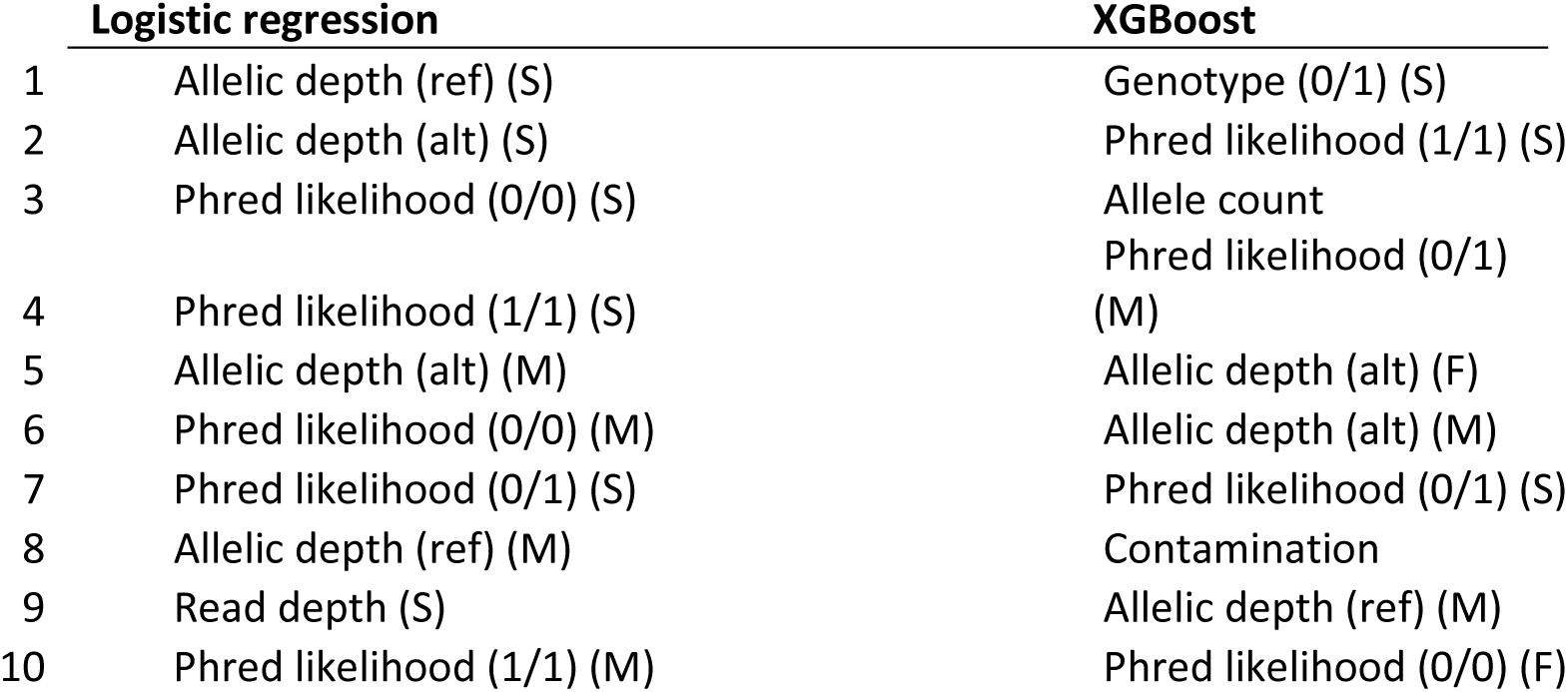
Top 10 most important features for machine learning methods in descending order. The parameters learned by the trained models (the coefficients of the hyperplanes separating the different classes for logistic regression; the degree of decision tree branching on each feature for XGBoost) can give an estimate of relative feature importance. Parentheses indicate the member of the trio (specimen, mother, father).

|  | Logistic regression | XGBoost |
| --- | --- | --- |
| 1 | Allelic depth (ref) (S) | Genotype (0/1) (S) |
| 2 | Allelic depth (alt) (S) | Phred likelihood (1/1) (S) |
| 3 | Phred likelihood (0/0) (S) | Allele count |
|  |  | Phred likelihood (0/1) |
| 4 | Phred likelihood (1/1) (S) | (M) |
| 5 | Allelic depth (alt) (M) | Allelic depth (alt) (F) |
| 6 | Phred likelihood (0/0) (M) | Allelic depth (alt) (M) |
| 7 | Phred likelihood (0/1) (S) | Phred likelihood (0/1) (S) |
| 8 | Allelic depth (ref) (M) | Contamination |
| 9 | Read depth (S) | Allelic depth (ref) (M) |
| 10 | Phred likelihood (1/1) (M) | Phred likelihood (0/0) (F) |

### Robustness of the genotype correction method

We explored the robustness of the XGBoost corrector performance to variations in the setup. First, since in practice, the MCC fraction is not known in advance, we investigated the robustness of the model to errors in MCC fraction estimate. We found that for the range of errors in the MCC estimate we obtained for the simulated specimens (Fig. S6), the decrease in performance is close to negligible (Fig. S9).

We then investigated the dependence of the genotype correction method performance on read coverage and found that for positions covered by more than 80 reads, which is typical coverage for exome sequencing, the genotype correction accuracy is close to maximal (Fig. S10). Finally, we investigated the effect of the choice of the reference genome on the performance of XGBoost. We tested the performance of the classifier with GRCh38 as the reference genome, and in settings where the classifier was trained on hg19-mapped data and tested on GRCh38-mapped data, and vice versa. As seen in Fig. S11, the machine learning approach is robust not only to the choice of the reference genome build, but also to changing the reference genome build between training and evaluation. Therefore, a model trained on one reference genome can be used to correct variants called with a different reference.

### Strelka2 genotype correction

To test the performance of the genotype-correction framework on a variant caller other than the GATK HaplotypeCaller, we repeated the tests with the VCF files obtained using the recently published caller Strelka2 [19] using its standard germline calling pipeline and the same VCF features as were available from GATK HaplotypeCaller. We observed the same trends, namely, that genotype correction increases the concordance between the calls on the “pure child” and the “contaminated specimen” trios (Fig. S12).

### Application to clinical miscarriage data

We applied the MCC estimation algorithm from section “Estimating the contamination fraction” to 33 abortus samples from a larger study aimed at finding genetic causes of spontaneous miscarriages of euploid fetuses (data to be published separately). In five of the 33 cases, the estimated MCC in the abortus exceeded 5%, with one case of 8% MCC and one of 40% MCC. We will focus on the one abortus sample with the 40% MCC (miscarried at 9 weeks) as one where MCC-aware variant correction can have the greatest impact (*cf*. Fig 2). We applied the recommended GATK hard filters [20] to the calls and further required that GQ be at least 30 for both parents. As a result, we obtained 65431 calls by MCC-naive GATK HaplotypeCaller before genotype correction, of which 11112 were genotype-corrected by the XGBoost (trained on the “virtual specimens”). As we expected, all but one corrections affected the heterozygous genotype call. Also not surprisingly, correction predominantly affected heterozygous variants with skewed allelic depths and favored the “elimination” of the allele with the lower allelic depth (Fig. S13).

Contamination-aware genotype correction had several effects, both on the global variant statistics and on the particular list of candidate variants that could explain the pregnancy loss. Genotype correction rectified the highly skewed ratio of heterozygous to homozygous variants called in the abortus and brought it in line with that for the uncontaminated parent samples. Thus, the parents’ heterozygous to homozygous call ratio, as computed by the GATK 3 VariantEval tool was 3.2-3.3, while for the MCC-naive abortus call set, it was 6.3. After genotype correction, that number for the abortus was brought down to 3.0, close to the parents’ value. Regarding the possible genetic explanation of pregnancy loss, the uncorrected call set contained six candidate compound heterozygote variants (Section S1; Table S1). Of these, genotype correction eliminated two variant calls, which reduced the number of candidate compound heterozygotes to four. Thus, in this case, MCC-aware variant (re-) calling eliminated potential false positive candidate variants.

## Discussion

As its costs are dropping, high-throughput sequencing is becoming increasingly routine in clinical and academic practice, and is gaining ground in prenatal diagnosis and in the diagnostic study of miscarried fetuses. While the bioinformatics toolkit for analyzing the sequencing data of individuals is well-developed and mature, it may encounter difficulties in the analysis of products of conception, which may be mixed with maternal tissue. We have shown that the accuracy of standard variant-calling pipelines does indeed degenerate as the maternal cell contamination increases, but that much of this decline can be alleviated by MCC-aware variant genotype correction. We have demonstrated that the MCC fraction can be readily estimated from trio sequencing data and then used to inform the genotype correction. We have further shown, using synthetic contaminated samples obtained from real trio data, that even a simple method based on Bayesian statistics does much to correct for the MCC-related loss of calling accuracy, and that the machine-learning methods trained on simulated data show even greater improvement, with XGBoost having the best performance.

We note that all methods discussed in this paper only correct the variants that have been discovered by a general-purpose variant caller (we used the popular GATK HaplotypeCaller, but any variant caller which outputs sufficiently rich information about variants into a VCF file should work). Therefore, MCC-aware genotype correction is a simple add-on to a standard bioinformatics pipeline, needing only the VCF file produced by it. The user can then compare the original VCF to the corrected VCF to see which genotypes had been changed. On the other hand, the reliance on standard variant calling implies that if MCC had caused the true fetal variant to be missed in all members of the trio, then that variant will not be called by the genotype-correction pipeline, either. We expect such cases to be rare.

Among published methods, CleanCall [21] applies its own probabilistic model to correct for sample contamination, starting with read-level information, and it has an option for doing so with a known contaminating sample (https://github.com/hyunminkang/cleancall). Since our method works with the output of a third-party variant caller, and comparing it to CleanCall cannot be decoupled from comparing CleanCall’s model to HaplotypeCaller’s, they are not directly comparable.

Nevertheless, we believe that in the case of maternal cell contamination with maternal sequence data available, our method is preferable, both because it is designed for that particular situation and because it allows the user to utilize the caller of their choice. To show the applicability of our method to real-world data, we applied it to in-house exome sequencing data for a spontaneously miscarried euploid embryo with 40% maternal contamination and demonstrated that it can correct for contamination artifacts. Currently, next-generation sequencing is applied to fetal DNA obtained by invasive methods which extract amniotic fluid or fetal or chorionic villi cells. Our work is aimed at the analysis of this sort of sequencing data, and we do not consider MCC fractions higher than 50%. As its costs drop, sequencing may become a practical option in non-invasive prenatal testing (NIPT) as well. NIPT analyzes fetal DNA circulating in maternal blood which constitutes a small fraction of the DNA sample obtained. Analyzing that sort of data would make MCC-aware variant calling a necessity, and may in fact give rise to more sophisticated and/or specialized algorithms, e.g., those that work directly with mapped reads.

### Genotype correction without the father’s information

We found that the XGBoost predictor, our most accurate method, does not benefit much from the inclusion of the father’s data. Since our experience with sample collection shows that obtaining the father’s DNA sample is often problematic, eliminating the need for father’s DNA should greatly expand the situations in which genotype correction can be applied. A difficulty lies in the MCC estimation algorithm’s reliance on the father’s genotype, but it can be circumvented, albeit at a possible loss of accuracy, either by adjusting the provided MCC-estimation algorithm or by using independent sources of information, such as the Short Tandem Repeat analysis. As a father-free analytical correction method, we provide a maximum-likelihood model that can be viewed as an approximation to the Restricted Bayesian model (Method S2).

## References

1. Tayoun ANA, Spinner NB, Rehm HL, Green RC, Bianchi DW. Prenatal DNA Sequencing: Clinical, Counseling, and Diagnostic Laboratory Considerations. Prenat Diagn. 2018;38(1):26–32.

2. Best S, Wou K, Vora N, Van der Veyver IB, Wapner R, Chitty LS. Promises, pitfalls and practicalities of prenatal whole exome sequencing. Prenat Diagn. 2018;38(1):10–19.

3. Stojilkovic-Mikic T, Mann K, Docherty Z, Ogilvie CM. Maternal cell contamination of prenatal samples assessed by QF-PCR genotyping. Prenat Diagn. 2005;25(1):79–83.

4. Weida J, Patil AS, Schubert FP, Vance G, Drendel H, Reese A, et al. Prevalence of maternal cell contamination in amniotic fluid samples. J Matern Fetal Neonatal Med. 2017;30(17):2133–2137.

5. Lamb AN, Rosenfeld JA, Coppinger J, Dodge ET, Dabell MP, Torchia BS, et al. Defining the impact of maternal cell contamination on the interpretation of prenatal microarray analysis. Genet Med. 2012;14(11):914–921.

6. Nagan N, Faulkner NE, Curtis C, Schrijver I. Laboratory Guidelines for Detection, Interpretation, and Reporting of Maternal Cell Contamination in Prenatal Analyses. J Mol Diagn. 2011;13(1):7–11.

7. DeBoever C, Aguirre M, Tanigawa Y, Spencer CCA, Poterba T, Bustamante CD, et al. Bayesian model comparison for rare variant association studies of multiple phenotypes. bioRxiv. 2018;doi: 10.1101/257162.

8. Degner JF, Marioni JC, Pai AA, Pickrell JK, Nkadori E, Gilad Y, et al. Effect of read-mapping biases on detecting allele-specific expression from RNA-sequencing data. Bioinformatics. 2009;25(24):3207–3212.

9. Jun G, Flickinger M, Hetrick KN, Romm JM, Doheny KF, Abecasis GR, et al. Detecting and estimating contamination of human DNA samples in sequencing and array-based genotype data. Am J Hum Genet. 2012;91(5):839–848.

10. Van der Auwera, G. (2014). Genotype refinement workflow. https://gatkforums.broadinstitute.org/gatk/discussion/4723/genotype-refinement-workflow.

11. GATK Team (2020). Genotype refinement workflow for germline short variants. https://gatk.broadinstitute.org/hc/en-us/articles/360035531432-Genotype-Refinement-workflow-for-germline-short-variants.

12. Chen T, Guestrin C. XGBoost: A scalable tree boosting system. 2016; Proceedings of the 22nd ACM SIGKDD, p. 785–794.

13. Zook JM, Catoe D, McDaniel J, Vang L, Spies N, Sidow A, et al. Extensive sequencing of seven human genomes to characterize benchmark reference materials. Scientific Data. 2016;3:160025.

14. Consortium, The 1000 Genomes Project. A global reference for human genetic variation. Nature. 2015;526(7571):68–74.

15. Jia Z, Fengbiao M, Wang L, Li M, Shi Y, Zhang B, et al. Whole-exome sequencing identifies a de novo mutation in TRPM4 involved in pleiotropic ventricular septal defect. Int J Clin Exp Pathol. 2017;10:5092–5104.

16. Corpas M, Valdivia-Granda W, Torres N, Greshake B, Coletta A, Knaus A, et al. Crowdsourced direct-to-consumer genomic analysis of a family quartet. BMC Genomics. 2015;16(1):910.

17. Jun G, Wing MK, Abecasis GR, Kang HM. An efficient and scalable analysis framework for variant extraction and refinement from population scale DNA sequence data. Genome Res. 2015; p. gr.176552.114. doi: 10.1101/gr.176552.114.

18. 1000 Genomes Project. GRCh38 Alignment README; 2015. https://github.com/igsr/1000Genomes_data_indexes/blob/master/data_collections/1000_genomes_project/README.1000genomes.GRCh38DH.alignment.

19. Kim S, Scheffler K, Halpern AL, Bekritsky MA, Noh E, Kallberg M, et al. Strelka2: fast and accurate calling of germline and somatic variants. Nat Methods. 2018;15(8):591.

20. Van der Auwera G. (howto) Apply hard filters to a call set; 2013. https://gatkforums.broadinstitute.org/gatk/discussion/2806/howto-apply-hard-filters-to-a-call-set.

21. Flickinger M, Jun G, Abecasis GR, Boehnke M, Kang HM. Correcting for Sample Contamination in Genotype Calling of DNA Sequence Data. Am J Hum Genet. 2015;97(2):284–290.

