## Supplemental Materials for "Accurate fetal variant calling in the presence of maternal cell contamination"

#### 1 Supplementary methods

##### S1 Variant prioritization

Variants that may be explanatory of pregnancy loss were prioritized as follows: candidate variants must be *de novo* variants or form compound heterozygotes. They are further prioritized based on the predicted deleteriousness of the variant (SIFT [1], PolyPhen2 [2], CADD [3]), population frequency (ExAc [4], gnomAD [5]), embryonic expressiont (GTEx [6], ExpressionAtlas [7]), clinical annotation (ClinVar [8], dbSNP [9]), prenatal lethality phenotype in mice (MGI [10]), and the affected proteins' intolerance to truncated variants [11]. The candidate mutations are manually selected based on this prioritization. The Ensembl Variant Effect Predictor [12] was additionally use in creating Table S1.

##### S2 Maximum likelihood-based genotype correction

As our simplest method, we consider a maximum likelihood-based approach that carries out a fetal genotype call adjustment based on a simple explicit mathematical model. To this end, we first observe that the disagreement between the true variants of the child and the called variants of the fetal specimen is in the vast majority of cases due to the mislabeling of homozygous positions in the fetus as heterozygous in the contaminated fetal specimen. This happens when the maternal genotype, and accordingly the mixture of fetal and maternal reads, is heterozygous.

Following this observation, we implement a fetal genotype correction procedure in which the candidates for readjustment are the positions where both the contaminated specimen's genotype and the maternal genotype are called heterozygous, say 01. We assume for simplicity that the maternal genotype 01 is known with high certainty (this can be ensured by filtering out positions with low genotype quality). We consider three hypotheses, that the child's genotype is 00, 01, or 11. We then determine the child's genotype using the maximum likelihood estimate based on the fetal allelic depths and the previously estimated MCC fraction $\alpha$. Specifically, given child's true genotype 00, 01 or 11 (and remembering that the maternal genotype is 01), the respective conditional probabilities of observing the allelic depths $AD0, AD1$ in the mixture are:

$$\begin{aligned}
\mathsf{P}(AD0, AD1|00) &= C(\tfrac{\alpha}{2})^{AD1}(1-\tfrac{\alpha}{2})^{AD0},\\
\mathsf{P}(AD0, AD1|01) &= C(\tfrac{1}{2})^{AD1}(\tfrac{1}{2})^{AD0},\\
\mathsf{P}(AD0, AD1|11) &= C(1-\tfrac{\alpha}{2})^{AD1}(\tfrac{\alpha}{2})^{AD0},
\end{aligned}$$

with the binomial coefficient $C = \frac{(AD0+AD1)!}{AD0!AD1!}$ Then, assuming for simplicity that the three possible fetal genotypes have the same prior probability, we just pick the genotype with the maximum probability.

#### S3 Bayesian correction

We describe a simple Bayesian model for estimating the probability of the child's genotype. The model assumes a single alternative allele, and ignores various subtleties present in practical variant calling pipelines, in particular estimation of genotype quality, filtering variants by genotype quality, elimination of repeated reads, etc. We make the following assumptions:

1. Let $AD0, AD1$ denote the allelic depths of the reference and alternative variants in the mother-fetus mixture, and $cGT, mGT, fGT$ denote the genotypes of child, mother and father, respectively. We assume that the joint probability of these variables factorizes as

$$\begin{aligned}
\mathsf{P}(AD0, AD1, cGT, mGT, fGT) = \mathsf{P}(&AD0, AD1|cGT, mGT)\\
&\times \mathsf{P}(cGT|mGT, fGT)\\
&\times \mathsf{P}(mGT)\cdot\mathsf{P}(fGT). \qquad (1)
\end{aligned}$$

   Here, the factors $\mathsf{P}(mGT)$ and $\mathsf{P}(fGT)$ are prior probabilities of mother's and father's genotypes, $\mathsf{P}(cGT|mGT, fGT)$ is the conditional probability describing Mendelian inheritance, and $\mathsf{P}(AD0, AD1|cGT, mGT)$ is the conditional probability describing observations of reads in the mixture for given genotypes of mother and child.

2. We assume that the prior probabilities $\mathsf{P}(mGT)$ and $\mathsf{P}(fGT)$ of mother's and father's genotypes $(00, 01$ or $11)$ are obtained from the phred-scaled likelihoods (code PL) of the respective VCF files.

3. The Mendelian conditional probabilities $\mathsf{P}(cGT|mGT, fGT)$ are given by

$$\mathsf{P}(c00|mGT, fGT) = \begin{cases} 1, & (m00, f00), \\ 1/2, & (m01, f00) \text{ or } (m00, f01), \\ 1/4, & (m01, f01), \\ 0, & \text{otherwise}, \end{cases}$$

$$\mathsf{P}(c11|mGT, fGT) = \begin{cases} 1, & (m11, f11), \\ 1/2, & (m01, f11) \text{ or } (m11, f01), \\ 1/4, & (m01, f01), \\ 0, & \text{otherwise}, \end{cases}$$

$$\mathsf{P}(c01|mGT, fGT) = \begin{cases} 1, & (m00, f11) \text{ or } (m11, f00), \\ 0, & (m00, f00) \text{ or } (m11, f11), \\ 1/2, & \text{otherwise}. \end{cases}$$

4. Each read of the mother-fetus mixture is assumed to belong to mother with probability equal to the previously estimated MCC fraction $\alpha$. All reads are assumed to be independent of each other. In addition, we assume the possibility of read errors: an allele can be called incorrectly (0 as 1, or 1 as 0) with probability $\epsilon$ (common for all reads). This gives

$$\begin{aligned}
&\mathsf{P}(AD0, AD1|m00, c00) = C(1-\epsilon)^{AD0}\epsilon^{AD1}, \\
&\mathsf{P}(AD0, AD1|m00, c01) = C(\tfrac{1+\alpha}{2} - \alpha\epsilon)^{AD0}(\tfrac{1-\alpha}{2} + \alpha\epsilon)^{AD1}, \\
&\mathsf{P}(AD0, AD1|m00, c11) = C(\alpha + (1-2\alpha)\epsilon)^{AD0}(1-\alpha-(1-2\alpha)\epsilon)^{AD1}, \\
&\mathsf{P}(AD0, AD1|m01, c00) = C(1-\tfrac{\alpha}{2}-(1-\alpha)\epsilon)^{AD0}(\tfrac{\alpha}{2}+(1-\alpha)\epsilon)^{AD1}, \\
&\mathsf{P}(AD0, AD1|m01, c01) = C(\tfrac{1}{2})^{AD0+AD1} \\
&\mathsf{P}(AD0, AD1|m01, c11) = C(\tfrac{\alpha}{2}+(1-\alpha)\epsilon)^{AD0}(1-\tfrac{\alpha}{2}-(1-\alpha)\epsilon)^{AD1} \\
&\mathsf{P}(AD0, AD1|m11, c00) = C(1-\alpha-(1-2\alpha)\epsilon)^{AD0}(\alpha+(1-2\alpha)\epsilon)^{AD1}, \\
&\mathsf{P}(AD0, AD1|m11, c01) = C(\tfrac{1-\alpha}{2}+\alpha\epsilon)^{AD0}(\tfrac{1+\alpha}{2}-\alpha\epsilon)^{AD1}, \\
&\mathsf{P}(AD0, AD1|m11, c11) = C\epsilon^{AD0}(1-\epsilon)^{AD1},
\end{aligned}$$

with the binomial coefficient $C = \frac{(AD0+AD1)!}{AD0!AD1!}$.

The above assumptions allow us to find the joint probability (1). The conditional probabilities $\mathsf{P}(cGT|AD0, AD1)$ are then determined by marginalizing it over $mGT, fGT$ :

$$\mathsf{P}(cGT|AD0, AD1) = Z^{-1} \sum_{mGT, fGT} \mathsf{P}(AD0, AD1, cGT, mGT, fGT),$$

where

$$Z = \sum_{mGT, fGT, cGT} \mathsf{P}(AD0, AD1, cGT, mGT, fGT).$$

Having found the probabilities $\mathsf{P}(cGT|AD0, AD1)$ for all possibilities of $cGT$, we choose the most probable genotype, i.e. $\text{argmax}_{cGT}\, \mathsf{P}(cGT|AD0, AD1)$.

### References


[1] R. Vaser et al. SIFT missense predictions for genomes. *Nature Protocols*, 11(1):1–9, January 2016.

[2] I. A. Adzhubei et al. A method and server for predicting damaging missense mutations. *Nature methods*, 7(4):248–249, April 2010.

[3] P. Rentzsch et al. CADD: predicting the deleteriousness of variants throughout the human genome. *Nucleic Acids Research*, 47(D1):D886–D894, January 2019.

[4] M. Lek et al. Analysis of protein-coding genetic variation in 60,706 humans. *Nature*, 536(7616):285–291, August 2016.

[5] K. J. Karczewski et al. Variation across 141,456 human exomes and genomes reveals the spectrum of loss-of-function intolerance across human protein-coding genes. *bioRxiv*, pp. 531210, January 2019.

[6] `https://gtexportal.org`.

[7] `https://www.ebi.ac.uk/gxa/`.

[8] `https://www.ncbi.nlm.nih.gov/clinvar/`.

[9] `https://www.ncbi.nlm.nih.gov/snp/`.

[10] Phenotypes and Mutant Alleles database at the Mouse Genome Informatics website, The Jackson Laboratory, Bar Harbor, Maine. World Wide Web . `http://www.informatics.jax.org`.

[11] C. A. Cassa et al. Estimating the selective effects of heterozygous protein-truncating variants from human exome data. *Nature Genetics*, 49:806, April 2017.

[12] W. McLaren et al. The Ensembl Variant Effect Predictor. *Genome Biology*, 17(1):122, June 2016.

#### 2 Supplementary figures

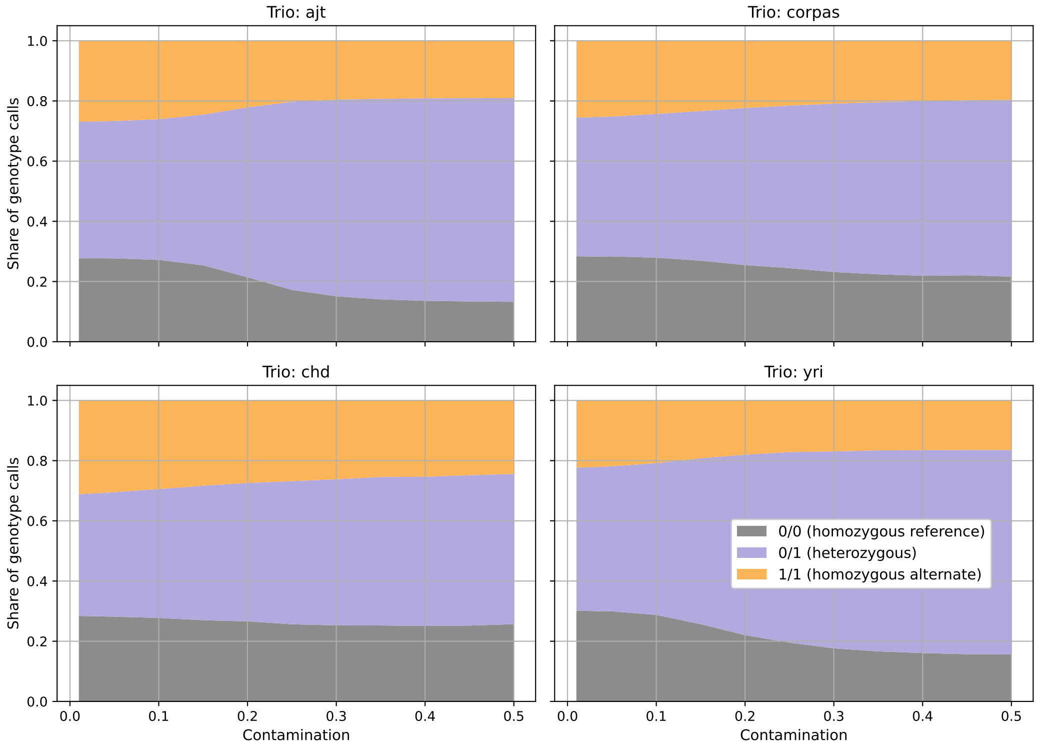


Figure S1: Visualization of the shares of specimen genotypes called as homozygous reference, heterozygous, and homozygous alternative by GATK at various MCC fractions.

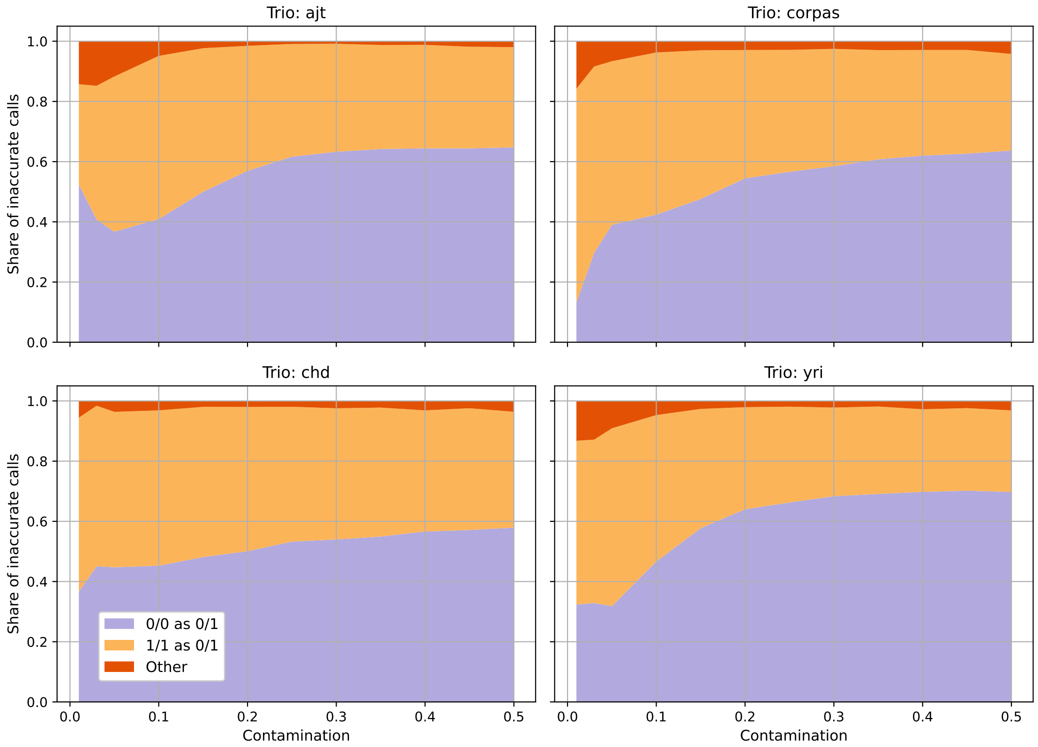


Figure S2: Visualization of the ratio of incorrectly called specimen genotypes (as compared to the ground truth) for various MCC fractions. The vast majority of incorrect calls are homozygous variants called as heterozygous.

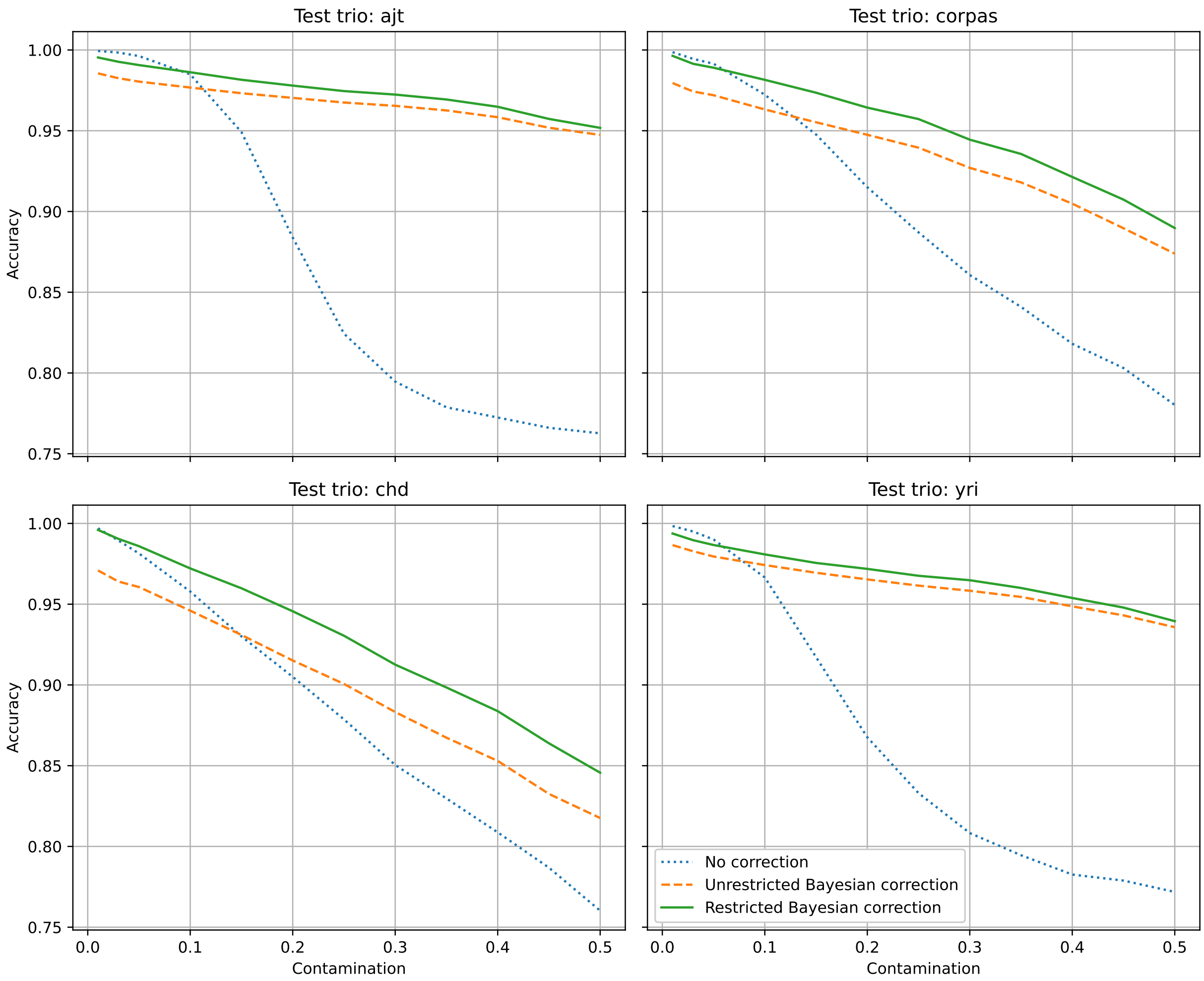


Figure S3: Accuracies of the restricted and unrestricted Bayesian correction methods at various MCC fractions.

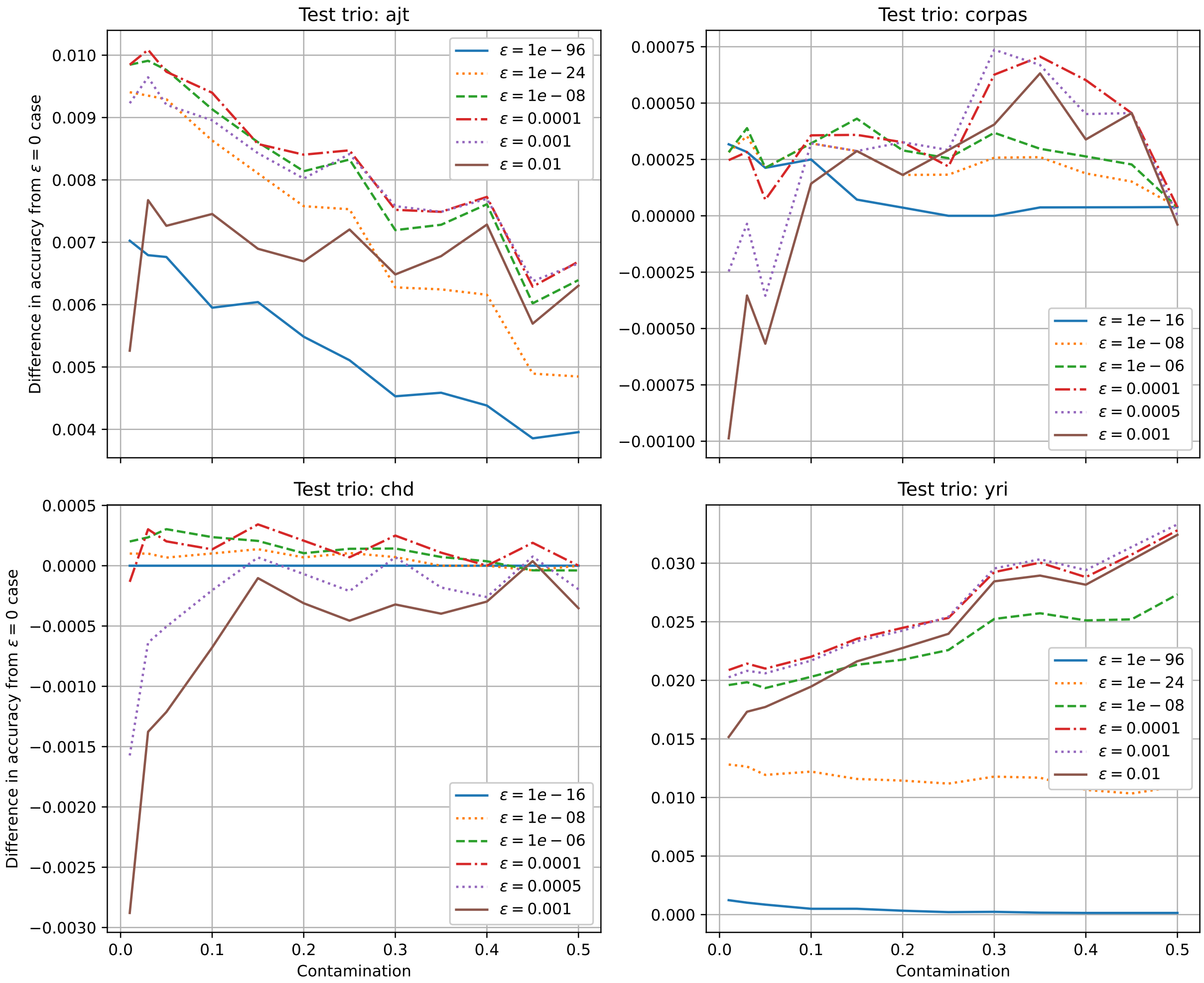


Figure S4: Difference between the accuracy of the unrestricted Bayesian correction for different values of $\epsilon$ from the accuracy of the $\epsilon = 0$ model. $\epsilon$ is the frequency of read errors assumed in the model.

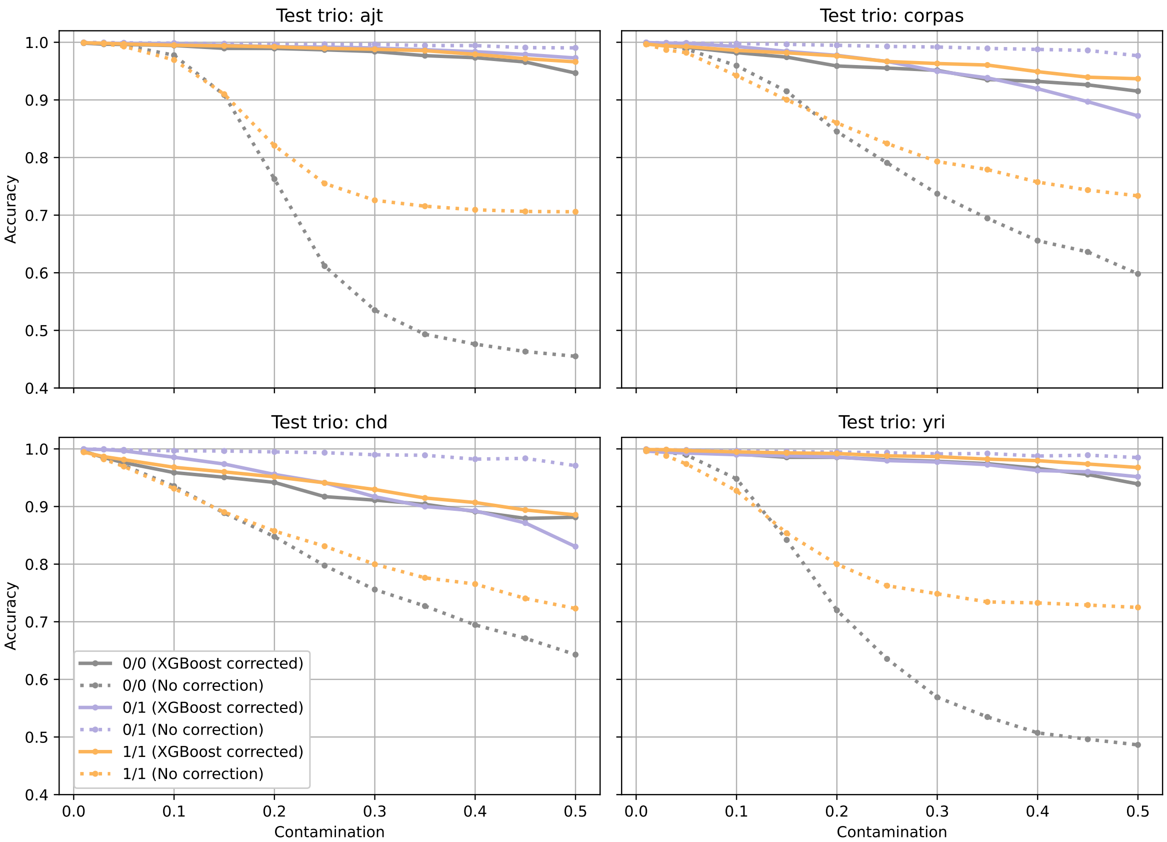


Figure S5: Accuracy before and after correcting with XGBoost visualized on a per-genotype basis.

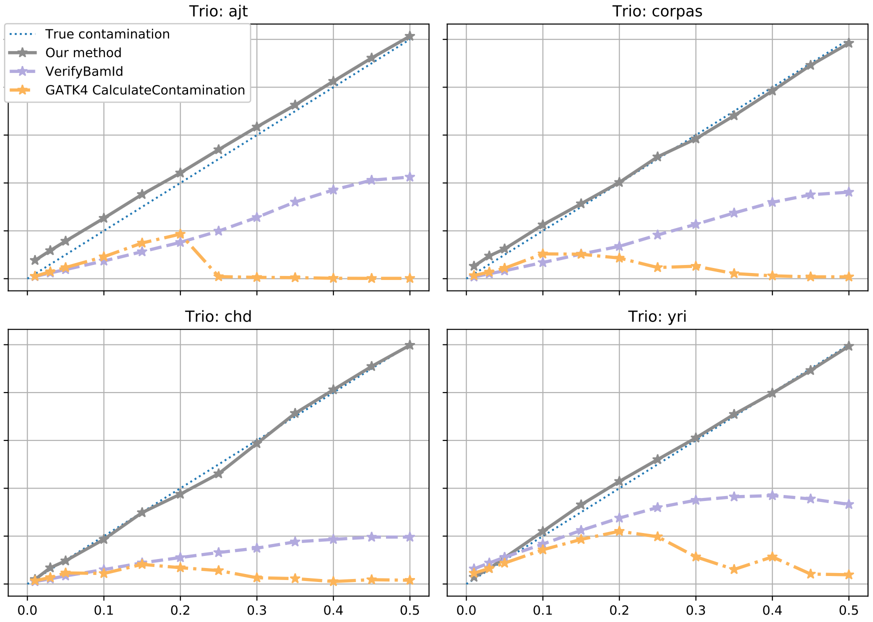


Figure S6: Estimates of the contamination fraction for virtual specimens generated from four real-world trios.

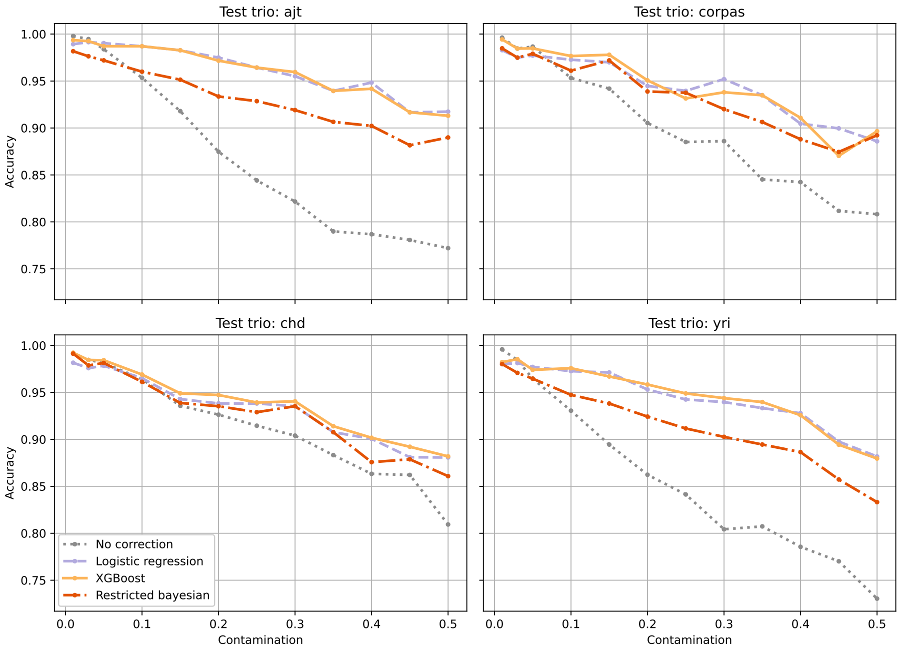


Figure S7: Performance of genotype correction on indels. The machine learning models were trained on variants that excluded indels, and tested on indel variants.

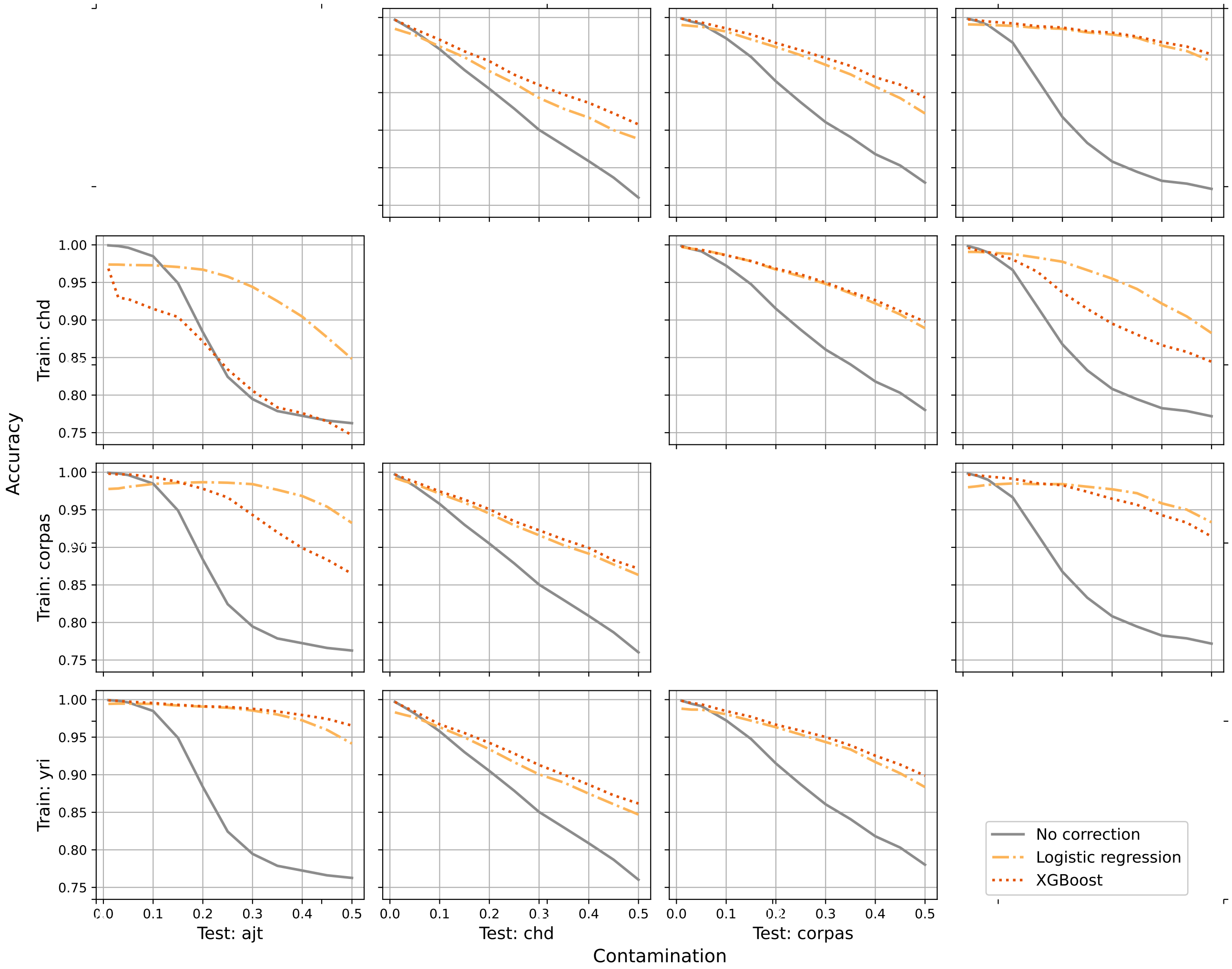


Figure S8: Accuracy of our methods as a function of contamination when trained and tested 'one versus one' on different pairs of trios. Each plot represents the case of training on one trio (varying along the vertical axis) and testing on another (varying on the horizontal axis).

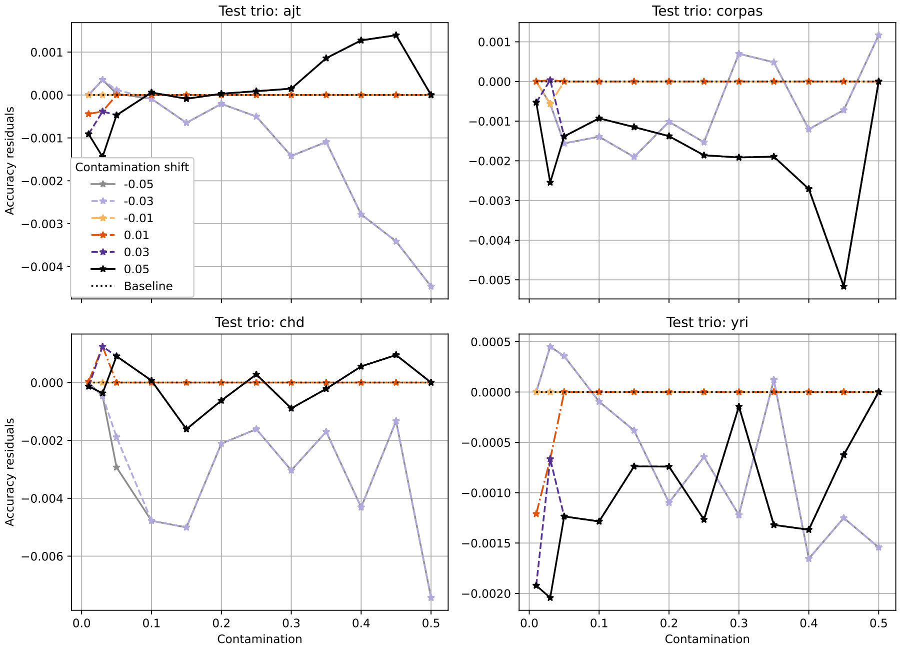


Figure S9: Change in accuracy of the XGBoost model as a function of contamination when the contamination estimate input to the algorithm is shifted from its true value by a constant amount (given by the labels on the plotted lines). Note the small range of variation in accuracy even for MCC estimation error of 5 percentage points.

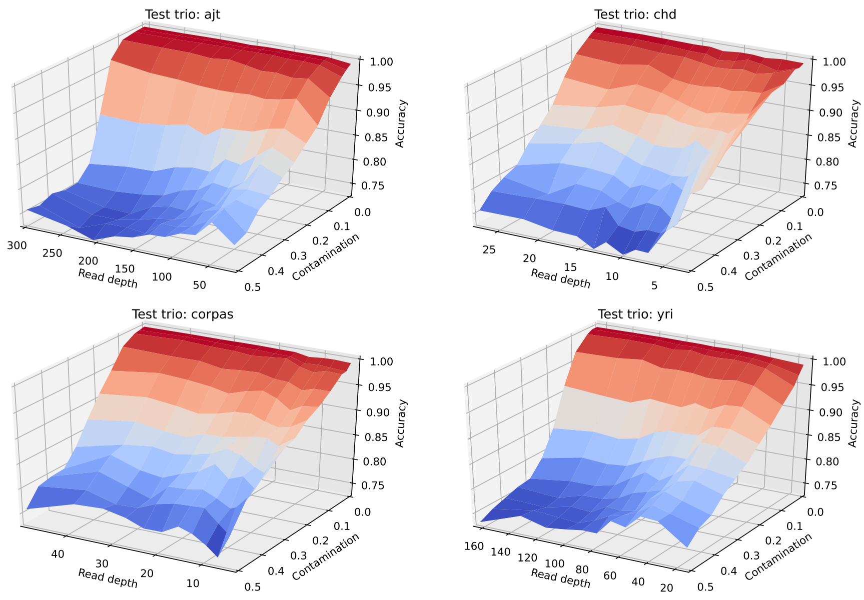


(a) No correction: full range

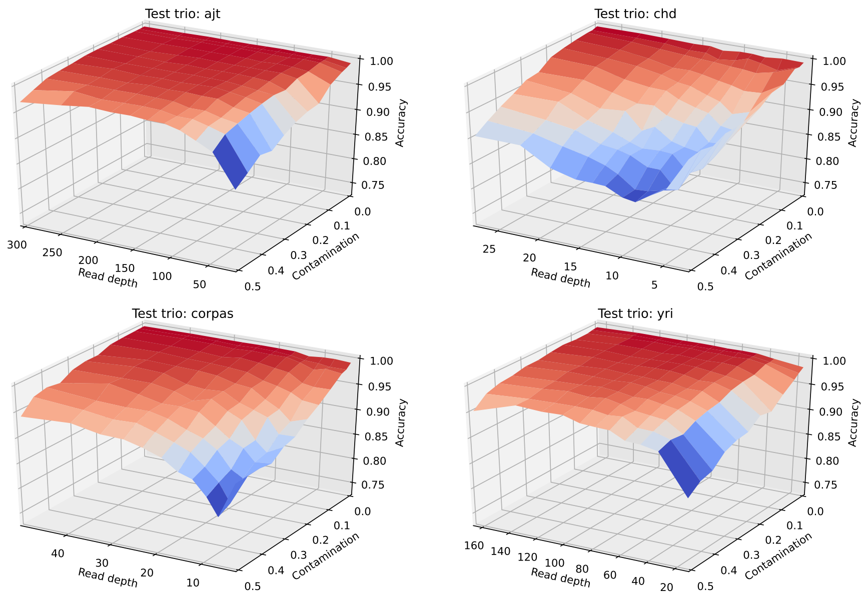


(b) XGBoost: full range

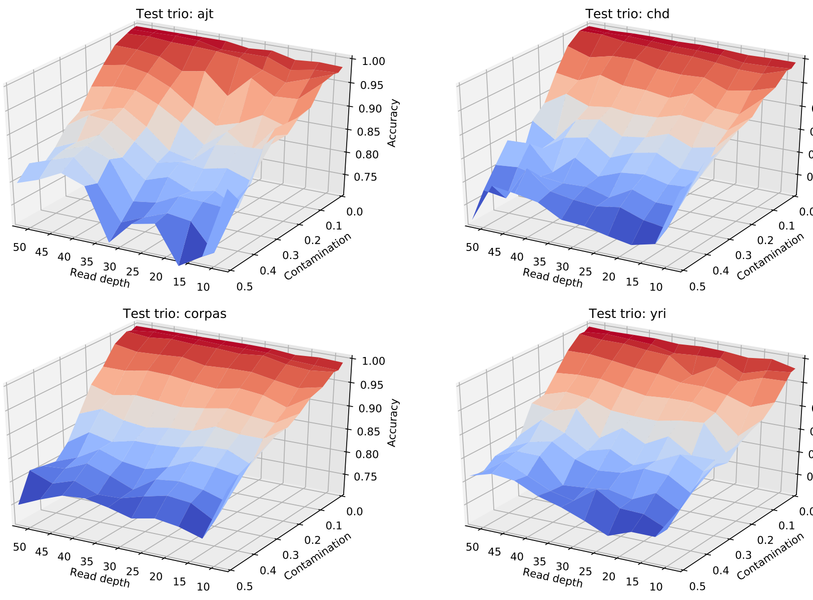


(c) No correction: 10x-50x coverage

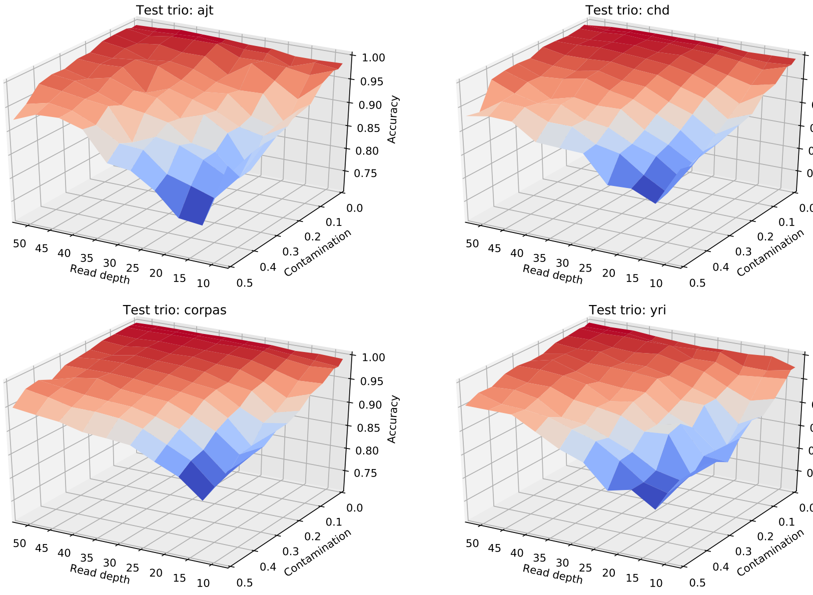


(d) XGBoost: 10x-50x coverage

Figure S10: Accuracy of our methods as a function of read depth and contamination. Subfigures (a) and (b) were obtained by stratifying positions into evenly distributed bins by read depth, and then calculating the accuracy for each bin. Subfigures (c) and (d) focus on 10x-50x read depth range, with evenly sized bins. Note that noise in regions of subfigures (c) and (d) can be attributed to the lower number of loci falling into these bins.

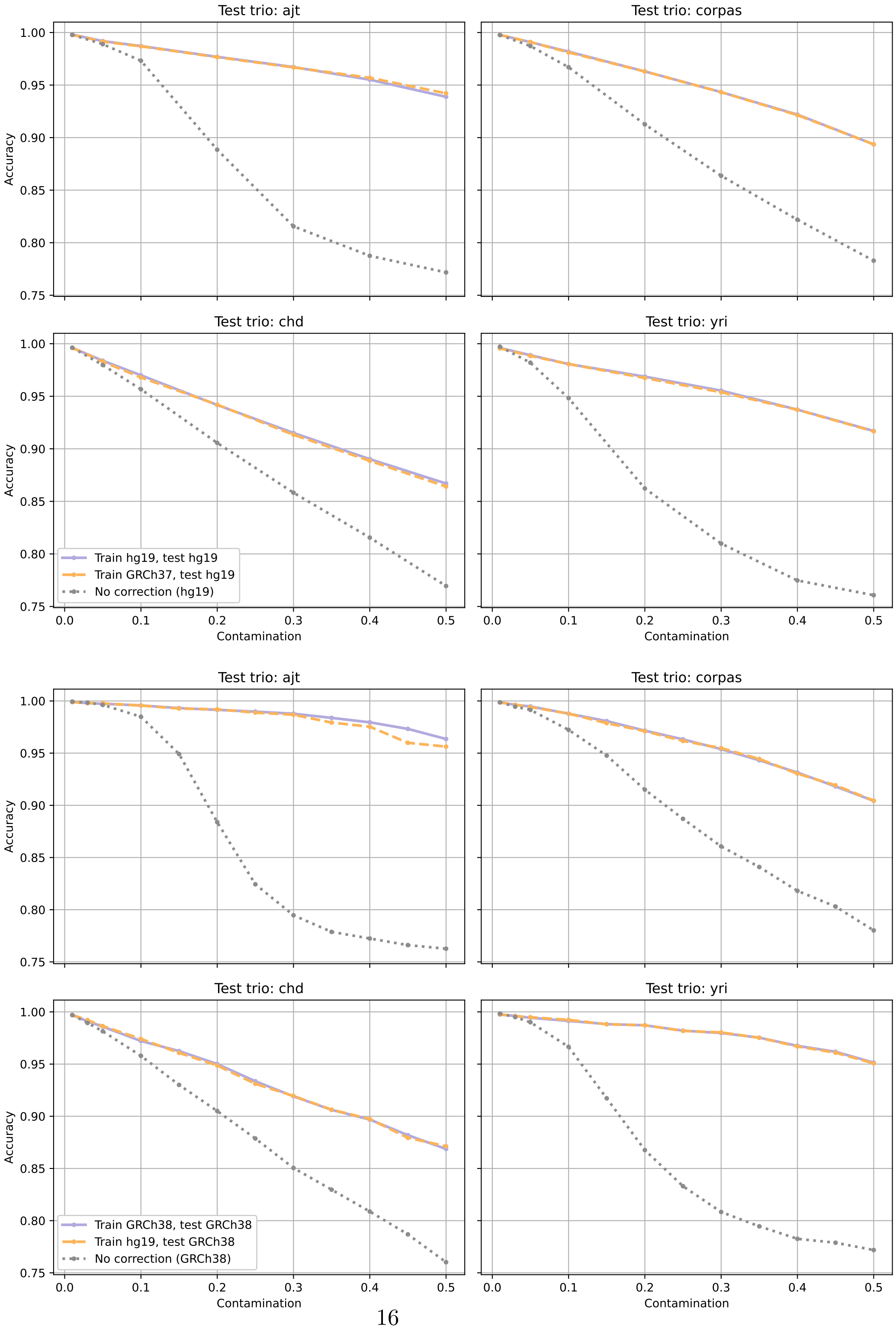




Figure S11: The effect of varying the training-data reference genome on the XGBoost model's correction accuracy. Top four: performance on the hg19-mapped test set. Bottom four: performance on the GRCh38-mapped test set.

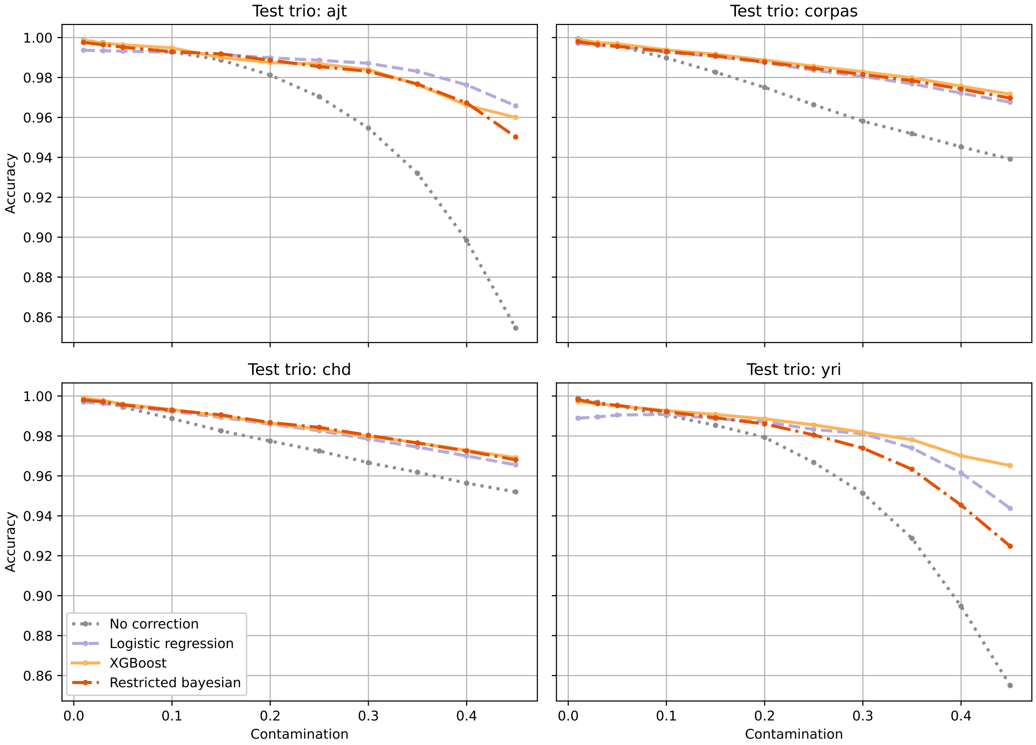


Figure S12: Accuracy of the genotype correction methods at various MCC fractions when applied to data called by the Strelka2 variant caller.

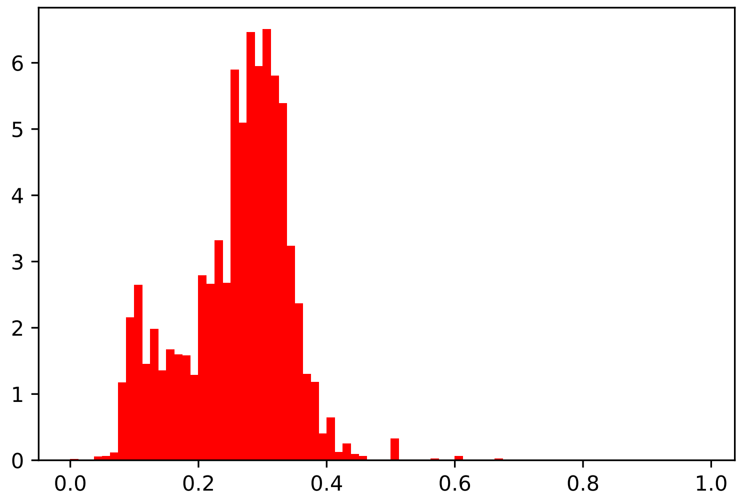


Figure S13: Distribution of the lower allelic fraction of the XGBoost-recalibrated variants for the miscarriage trios.

### 3 Supplementary table

| chrom | position | gene | transcript change | aa change | alt allele frac | rs id |
|---|---|---|---|---|---|---|
| chr1 | 22160001 | HSPG2 | NM_005529.7:c.10937G>A | p.(Arg36476His) 98 | 0.2 | rs112062179 |
| *** chr1 | 22215199 | HSPG2 | ENST00000374673.3:c.17C>A† | p.Pro6Gln | 0.158 | rs202082510 |
| chr1 | 186277040 | PRG4 | NM_005807.5:c.2189C>T | p.(Pro730Leu) | 0.28 | rs201317498 |
| chr1 | 186277625 | PRG4 | NM_005807.5:c.2774G>A | p.(Arg925His) | 0.438 | rs142388523 |
| chr1 | 186276404 | PRG4 | NM_005807.5:c.1553C>T | p.(Thr518Ile) | 0.293 | None |
| chr12 | 129178533 | TMEM132C | NM_001136103.2:c.1609C>A | p.(Leu537Ile) | 0.344s | rs191481498 |
| *** chr12 | 129180617 | TMEM132C | NM_001136103.2:c.1898T>C | p.(Val633Ala) | 0.098 | None |
| chr17 | 73258028 | MRPS7 | NM_015971.4:c.47C>T | p.(Ala16Val) | 0.409 | rs148590649 |
| chr17 | 73258570 | MRPS7 | NM_015971.4:c.84-8C>T | splice region variant‡ | 0.207 | rs200570062 |
| chr2 | 173832019 | RAPGEF4 | NM_007023.4:c.851A>G | p.(Tyr284Cys) | 0.246 | rs202063054 |
| chr2 | 173881919 | RAPGEF4 | NM_007023.4:c.1916C>T | p.(Ala639Val) | 0.333 | rs150495482 |
| chr6 | 152469204 | SYNE1 | NM_182961.4:c.24952C>T | p.(Leu8318Phe) | 0.317 | rs141716975 |
| chr6 | 152757224 | SYNE1 | NM_182961.4:c.4162C>T | p.(Arg1388Trp) | 0.382 | rs34028822 |
| chr6 | 152774753 | SYNE1 | NM_182961.4:c.2995G>A | p.(Glu999Lys) | 0.306 | rs148346599 |

Table S1: Candidate variants causative for pregnancy loss in PG16 (the genomic coordinates are relative to the hg19 assembly). Asterisks mark variants that were eliminated by genotype correction. Variants were annotated with the Ensembl VEP tool [12].

†The chr1:22215199 G>T leads to an amino acid change in the Ensembl transcript ENST00000374673.3, whereas it is an intron variant in the RefSeq transcripts.

‡The chr17:73258570 C>T is missense in the Ensembl transcript ENST00000579002.1 (c.163C>T; p.(Pro55Ser)).
